# Highly efficient site-directed gene insertion in primary human natural killer cells using homologous recombination and CRISPaint delivered by AAV

**DOI:** 10.1101/743377

**Authors:** Meisam Naeimi Kararoudi, Shibi Likhite, Ezgi Elmas, Maura Schwartz, Kathrin Meyer, Dean A. Lee

## Abstract

Human peripheral blood natural killer (NK) cells have strong antitumor activity and have been used successfully in several clinical trials. Modifying NK cells with a chimeric antigen receptor (CAR) can improve their targeting and increase specificity. However, genetic modification of NK cells has been challenging due to the high expression of innate sensing mechanisms for viral nucleic acids. Recently, we described an efficient vector-free method using Cas9/ribonucleoprotein complexes for gene deletion in NK cells. Here, we combined this approach with single-stranded (ss) or self-complementary (sc) Adeno-associated virus (AAV)-mediated gene delivery for gene insertion into a user-defined locus using homology repair (HR) and non-homologous directed CRISPR-assisted insertion tagging (CRISPaint) approaches. Using these approaches, we identified scAAV6 as the superior serotype for successful generation of stable mCherry-expressing primary NK cells (up to 89%). To maximize transgene packaging in HR-directed gene insertion, we identified minimum optimal homology arm lengths of 300bp for the flanking region of the Cas9-targeting site. Lastly, we demonstrate that mCherry positive NK cells can be expanded to large numbers using feeder cells expressing membrane-bound IL-21. This efficient method for site-directed insertion of genetic material into NK cells has broad potential for basic discovery and therapeutic applications for primary NK cells.

## Introduction

Gene modification of NK cells using viral or non-viral vectors has been challenging due to robust foreign DNA- and RNA-sensing mechanisms, and hence limits the efficiency of gene delivery methods into NK cells. To overcome this limitation, we have recently developed a new method to electroporate Cas9/ribonucleoprotein complexes (Cas9/RNP) directly into human primary NK cells. This method introduces a double-strand break (DSB) in the genome of NK cells, which results in successful gene knock-out and enhanced antitumor activity^1^. Cas9 protein is preferable to mRNA delivery due to its fast action and clearance^2^. After this initial success in gene silencing, we thought to further develop the method for gene insertion. Following the action of Cas9 of introducing a DSB, two independent pathways may be utilized to repair the damage, known as homologous recombination (HR) and homology-independent repair. In the presence of a DNA template encoding a gene of interest, the exogenous gene can be integrated into the Cas9-targeting site using either repair mechanisms^3^. There are several ways to provide such a DNA template, including viral and non-viral methods. In non-viral approaches, the single-stranded or double-stranded DNA template is typically electroporated along with Cas9/RNP^4^. For viral gene delivery, adeno-associated viruses (AAV) were used safely in clinical trials and are effective as vectors for sensitive primary immune cells, including T-cells^5^. Transcripts that are delivered via AAV vectors can be packaged as a linear single-stranded (ss) DNA with a length of approximately 4.7 kb (ssAAV) or linear self-complementary (sc) DNA (scAAV). scAAV contains a mutated ITR which helps to bypass rate-limiting steps of second strand generation for converting ssDNA into double-stranded (ds)DNA^6^. However, as a consequence, the scAAV has only half the packaging capacity compared to ssAAV and hence is not suitable for larger transgenes^6^. Therefore, we designed and tested both ssAAV and scAAV for DNA template delivery into NK cells.

Stable gene integration through HR machinery depends on providing the transgenes with optimal homology arms for the flanking region of the DSB^7-10^. To find the most optimal length of homology arms and optimize packaging capacity of a transgene into ssAAV and scAAV, we designed 30bp, 300bp, 500bp and 800-1000 bps of HAs for the right and left side of Cas9-targeting site. Since designing homology arms is a time-consuming procedure and requires multiple optimizations, we also investigated the CRISPaint approach, a homology-independent method for gene insertion or tagging. In this method, the same Cas9 targeting site, including crRNA and PAM sequences, is provided in the DNA template encoding the gene of interest. Upon introduction of the Cas9 complex, both template and genomic DNA will be cut simultaneously. As a result, the CRISPaint template will be presented as a linearized double-stranded DNA which can be integrated through non-homology repair machinery^4, 11^. With both methods we successfully generated highly efficient and stable transgene-modified NK cells using mCherry as a proof of concept.

## Methods

### Human NK Cell Purification and Expansion

NK cells were purified as previously described^1^. Briefly, NK cells were isolated from PBMC using RosetteSep™ Human NK Cell Enrichment Cocktail (Figure 1). Purified NK cells were phenotyped using flow cytometry as >90% CD3-negative/CD56-positive population (Figure 2). These cells were stimulated with irradiated mbIL21-expressing K562 feeder cells at a ratio of 2:1 (feeder:NK) at the day of purification (Figure 1) ^12^. The stimulated cells were cultured for 7 days in the serum-free AIM-V/ICSR expansion medium containing 100 IU/mL of IL-2.

**Figure 1.**
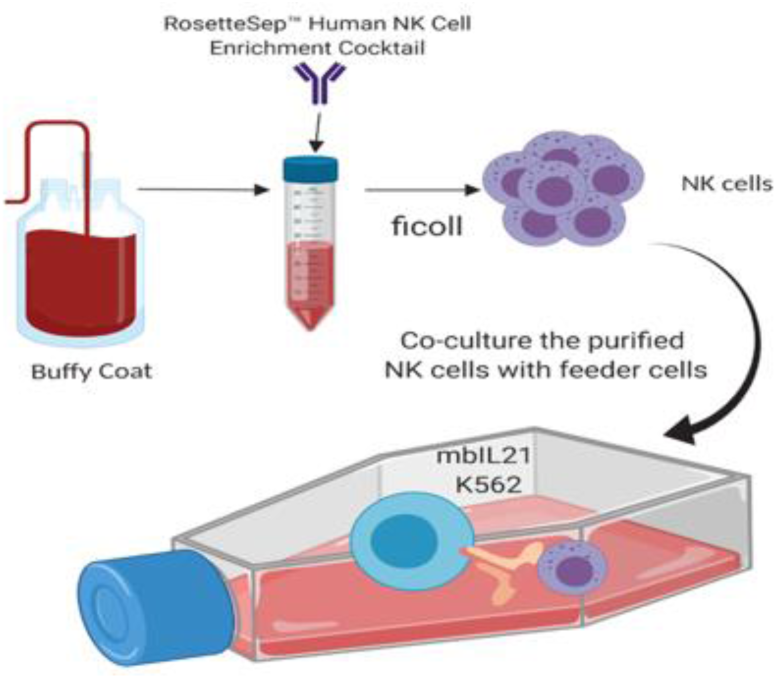
Scheme of NK isolation and expansion. Primary human NK cells were isolated from buffy coats from healthy donor and expanded using mbIL21 K562 for 7 days.

**Figure 2.**
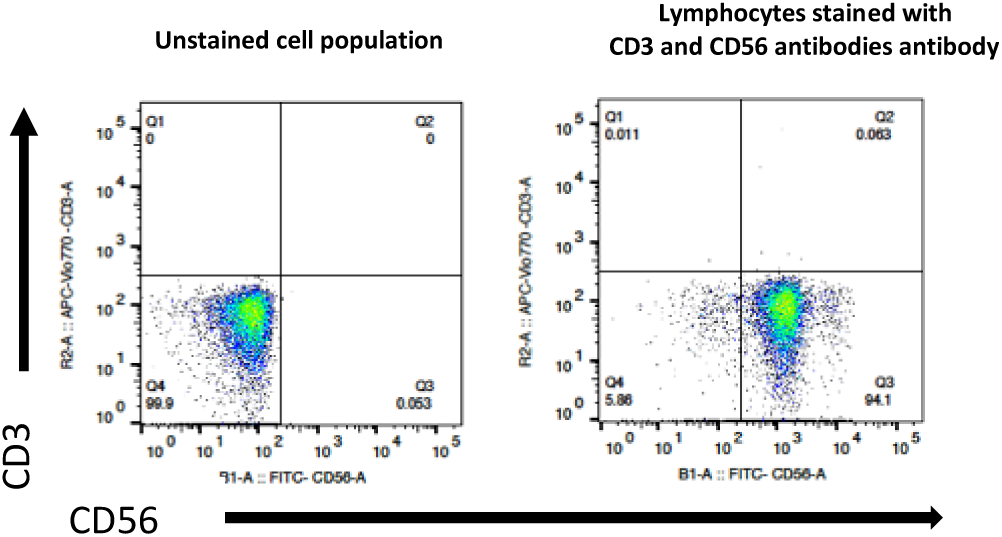
Flow cytometry result of isolated NK cells shows the purity of NK cells. The cells isolated from buffy coat were evaluated for T-cell contamination by staining for CD3 and CD56. Based on the flow cytometry results, the isolated lymphocytes were >90% CD3-negative, CD56-positive.

### Targeting Genomic safe harbors for gene insertion

Genomic safe harbors (GSHs) are sites in the genome which can be modified with no change in the normal function of the host cell and allow adequate expression of the transgene. For our proof of concept study for gene insertion in NK cells, we chose the *adeno-associated virus site 1 (AAVS1)*, which is one of the GSHs and an exemplary locus within the *phosphatase 1 regulatory subunit 12C (PPP1R12C)* gene. This locus has been successfully used for directed gene insertion into several cell types^3, 13^. First, we evaluated the chromatin accessibility of AAVS1 in naïve and expanded NK cells by ATAC-seq assay and showed no difference between the naïve and IL-21 expanded cells (Figure 3). AAVS1 was targeted using one gRNA (crRNA: 5’GGGGCCACTAGGGACAGGAT) via electroporation of Cas9/RNP into expanded NK cells as described before^1^.

**Figure 3.**
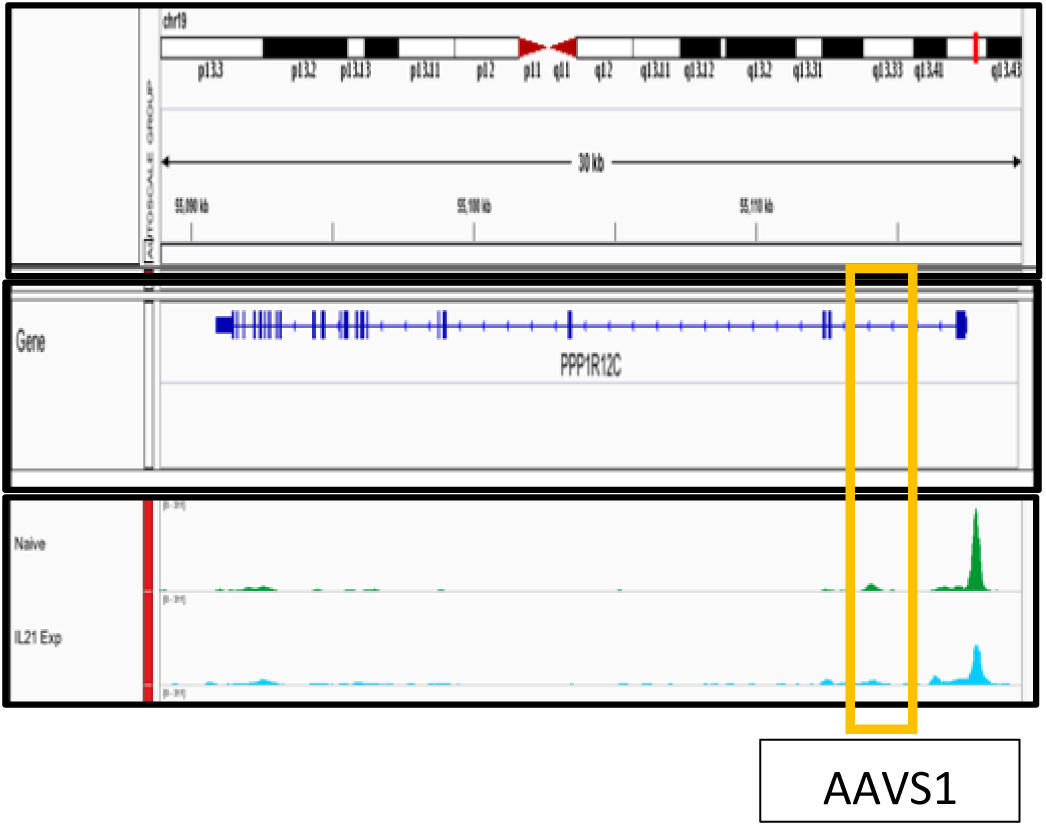
ATAC-seq analysis shows that AAVS1 in NK cells has suitable chromatin accessibility for gene insertion. ATAC-seq peaks representing open chromatin regions. The data shows AAVS1 has similar chromatin accessibility in naïve and expanded day 7 NK cells.

Briefly, 3 × 10^6^ expanded NK cells were harvested and washed twice with 13ml of PBS followed by centrifugation for 7 minutes at 300g and aspiration of PBS. The cell pellet was resuspended in 20ul of P3 Primary Cell 4D-Nucleofector Solution. 5ul of pre-complexed Cas9/RNP (Alt-R® CRISPR-Cas9 crRNA, Alt-R® CRISPR-Cas9 tracrRNA and Alt-R® S.p. HiFi Cas9 Nuclease V3) (Integrated DNA Technologies, Inc., Coralville, Iowa), targeting AAVS1 and 1ul of 100uM electroporation enhancer (Alt-R® Cas9 Electroporation Enhancer) were added to the cell suspension. The total volume of 26ul of CRISPR reaction was transferred into 4D-Nucleofector^™^ 16-well Strip and electroporated using program EN-138. After electroporation, the cells were transferred into 2ml of serum-free media containing 100IU of IL-2 in a 12 well plate (Figure 4) and incubated at 37 degrees 5% CO2 pressure.

**Figure 4.**
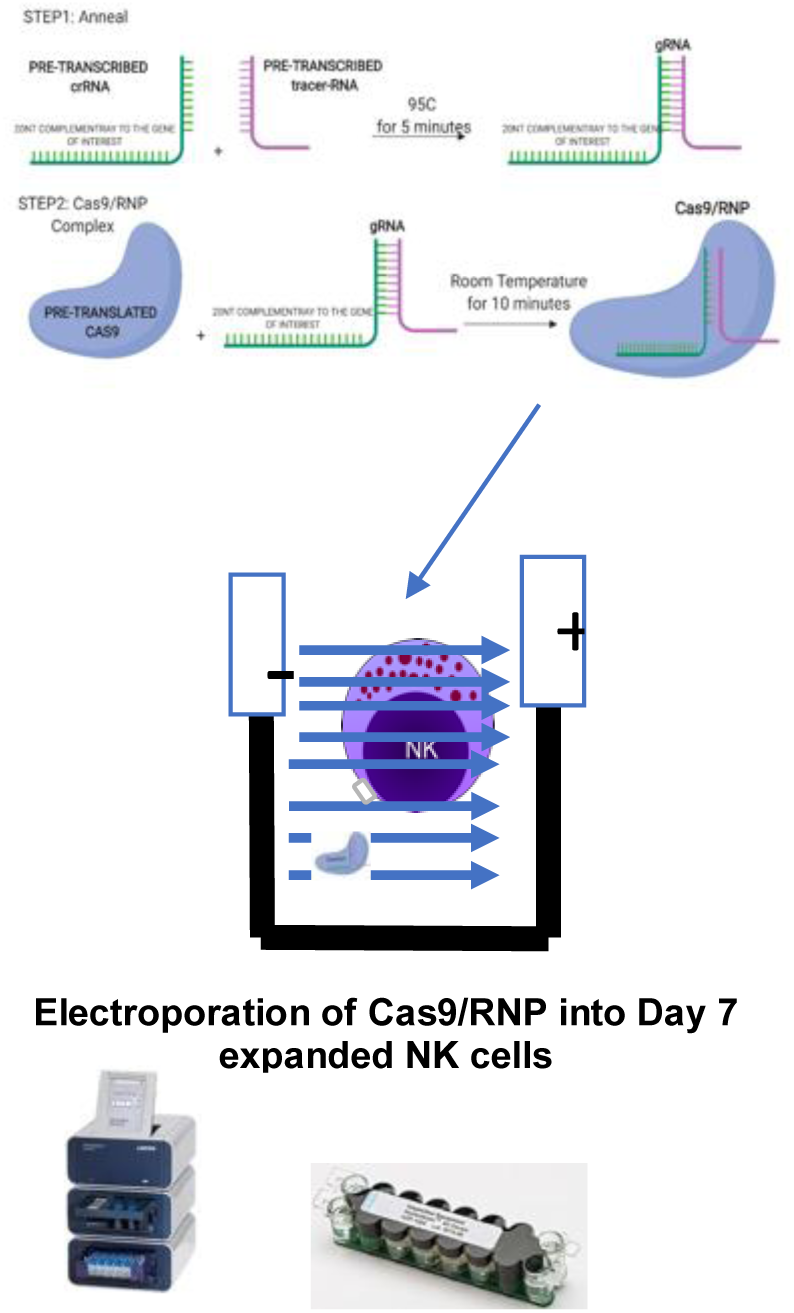
Production and electroporation of Cas9/RNP to target the gene of interest in human primary NK cells. To make the gRNA, pre-transcribed crRNA and Tracer-RNA were annealed at 95C for 5 minutes. Then pre-translated Cas9 endonuclease protein was mixed with gRNA at room temperature for 10 minutes. The Cas9/RNP and electroporation enhancer were added to NK cells. The mix then was transferred into Lonza4D nucleofection cuvette. The cells were electroporated with EN-138 program using Lonza 4D.

After 48 hours, NK cell DNA was isolated for detection of Insertions deletions (Indels) in CRISPR edited NK cells. The region flanking the Cas9 targeting site was PCR amplified and the amplicons were Sanger sequenced. Inference of CRISPR Edits (ICE) was used to analyze the frequency of Indels (Figure 5) ^14^. The ICE results showed that more than 85% of CRISPR modified NK cells had at least one indel at the AAVS1 Cas9-targeting site. To ensure that genome modifications at this locus did not interfere with the ability to target cancer cells, we studied cytotoxicity of AAVS1-KO NK cells against Kasumi, an AML cancer cell line. Using a Calcein AM assay, we observed no difference between wild type and CRISPR modified NK cells^1^ in their killing ability (Figure 6).

**Figure 5.**
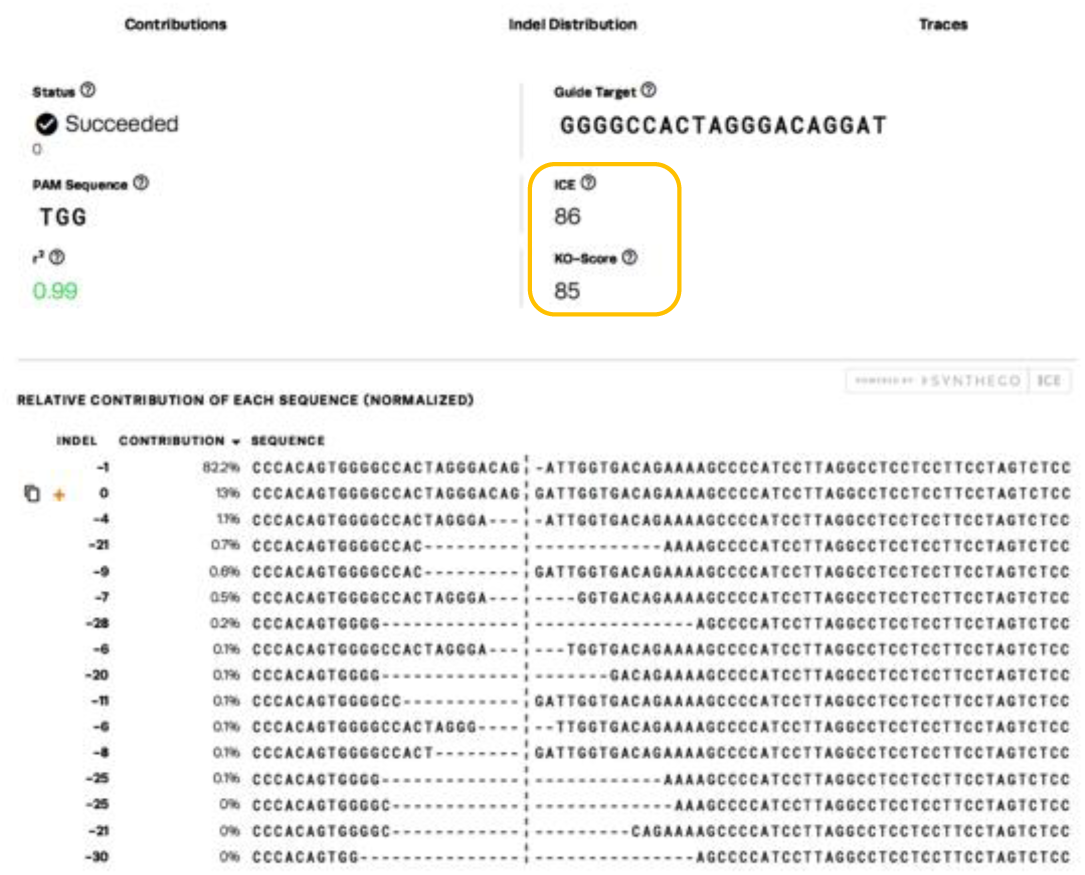
The ICE analysis shows successful gene (AAVS1) targeting in NK cells using Cas9/RNP. The ICE uses sanger sequencing data to produce quantitative analysis of CRISPR editing. Using this method, we showed that more than 85% of NK cells were successfully targeted at AAVS1 locus.

**Figure 6.**
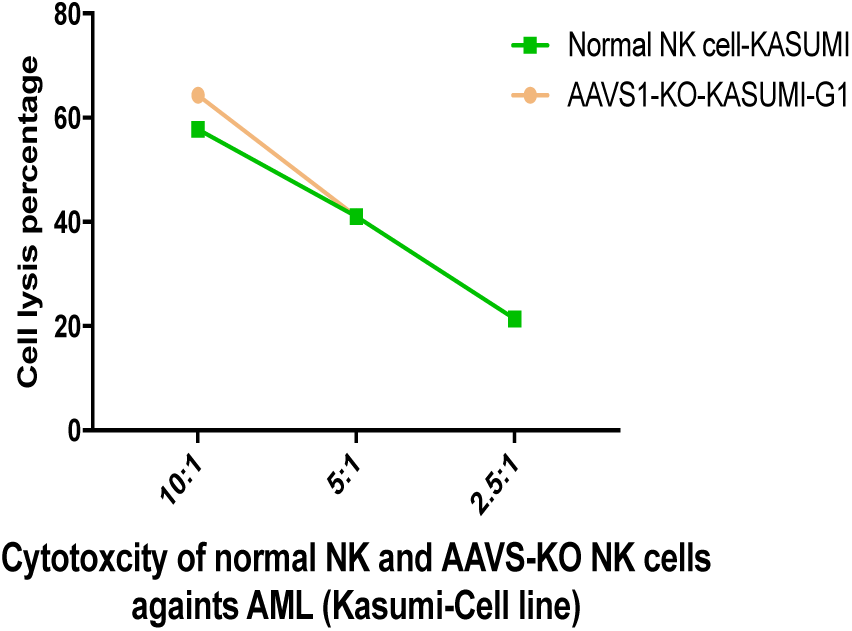
Knocking-out AAVS1 does not alter NK cells cytotoxcicty. Calcein-AM cytotoxicity assay showed no difference in NK cell-mediated killing against AML cell line by AAVS1-KO and wildtype NK cells. This demonstrates the safety of targeting AAVS1 locus in NK cells.

### Testing of various Adeno-Associated Viral Vector serotypes for DNA template delivery

To determine the best serotype of AAV for transduction of primary NK cells and to provide the highest number of DNA template encoding gene of interest into NK cells, we transduced the cells with several serotypes of AAVs including AAV4, AAV6, AAV8 and AAV9 encoding GFP at a multiplicity of infection (MOI) of 300K. We saw that NK cells transduced with AAV6 had the highest expression level of GFP detected by flow cytometry. Importantly, the AAV6 viral genome could be detected up to 48 hours post-transduction by qPCR analysis, which is a critical time for the endonuclease function of Cas9 protein (Figure 7).

**Figure 7.**
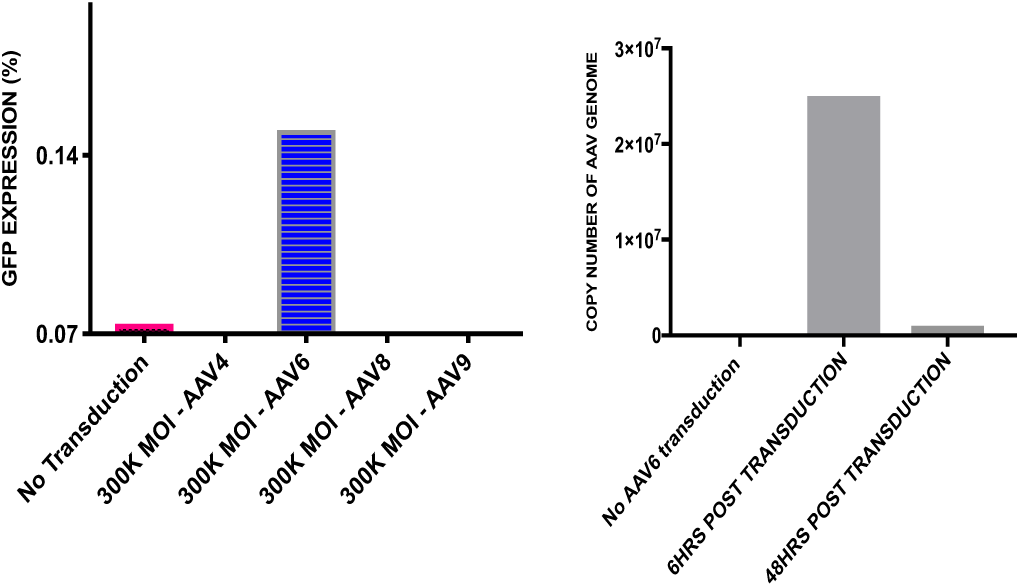
Identifying the best serotype of AAV to transduce human primary NK cells. Among 4 different serotypes of AAVs, only NK cells that were transduced with AAV6 showed expression of GFP. AAV6 viral genome could be detected after 48 hours post transduction of NK cells at MOI of 300K.

### Designing HR and CRISPaint gene delivery contructs

After successful proof of concept that the AAVS1 locus can be modified in NK cells without altering their ability to target cancer cells, we evaluated AAV mediated delivery of the mCherry transgene for gene insertion at AAVS1. For HR-directed gene insertion, DNA-encoding mCherry with homology arms (HA) to the flanking region of Cas9-targeting site were cloned into the backbone of single-stranded or self-complementary AAV vector ^15^. We designed 30bp, 300bp, 500bp and 1000bp of HA for the right and 30bp, 300bp, 500bp and 800bp for the left HA (Figure 8). For the CRISPaint DNA templates, we incorporated single (PAMg) or double (PAMgPAMg) Cas9-targeting sequences around the mCherry transgene but within the ITRs. Therefore, Cas9 could simultaneously cut gDNA and the CRISPaint DNA template, enabling integration at the genomic DSB (Figure 9). To ensure the accuracy of the designed HR and CRISPaint DNA templates before packaging the transgenes into AAV6, we co-electroporated the circular DNA encoding mCherry into HEK293 cells with Cas9/RNP targeting AAVS1. The results showed that both HR and CRISPaint DNAs were successfully integrated at the genomic DSB and mCherry was efficiently expressed (Figure 10). The production of both single-stranded and self-complementary AAV6 viral particles was done as previously described ^15^.

**Figure 8.**
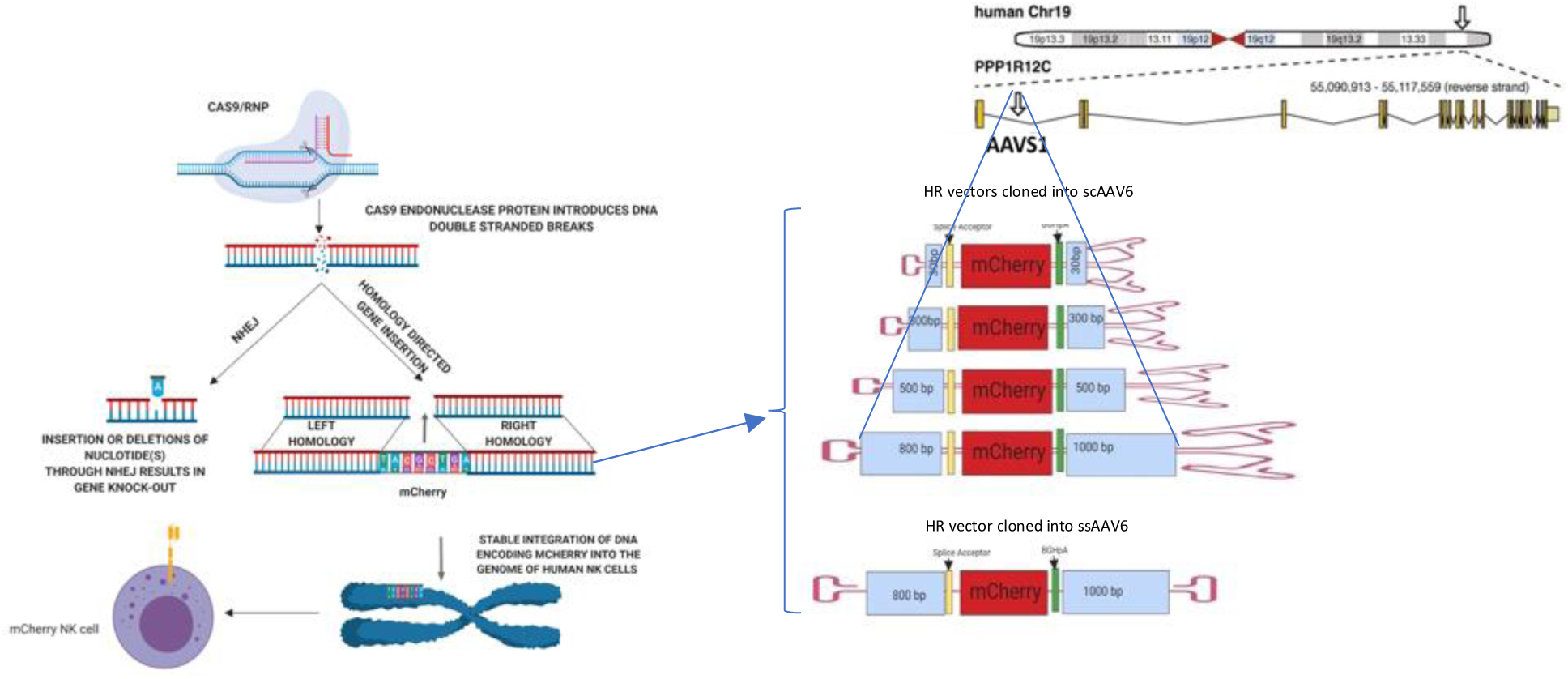
HR-directed gene insertion requires optimal length of HAs. Non-Homologues End Joining (NHEJ) DNA repair machinery is activated following Cas9 DSB which results in gene knock-out. The Homology Repair (HR) DNA repair machinery is the pathway which its activation results in gene insertion in presence of a DNA template with optimum homology arms for Cas9-targeting site (AAVS1). We designed 30bp, 300bp, 500bp and 800bp of HA for the left HA and 30bp, 300bp, 500bp and 1000bp for the right HA. The HR templates were cloned within ITRs of ssAAV6 or scAAV6 vectors.

**Figure 9.**
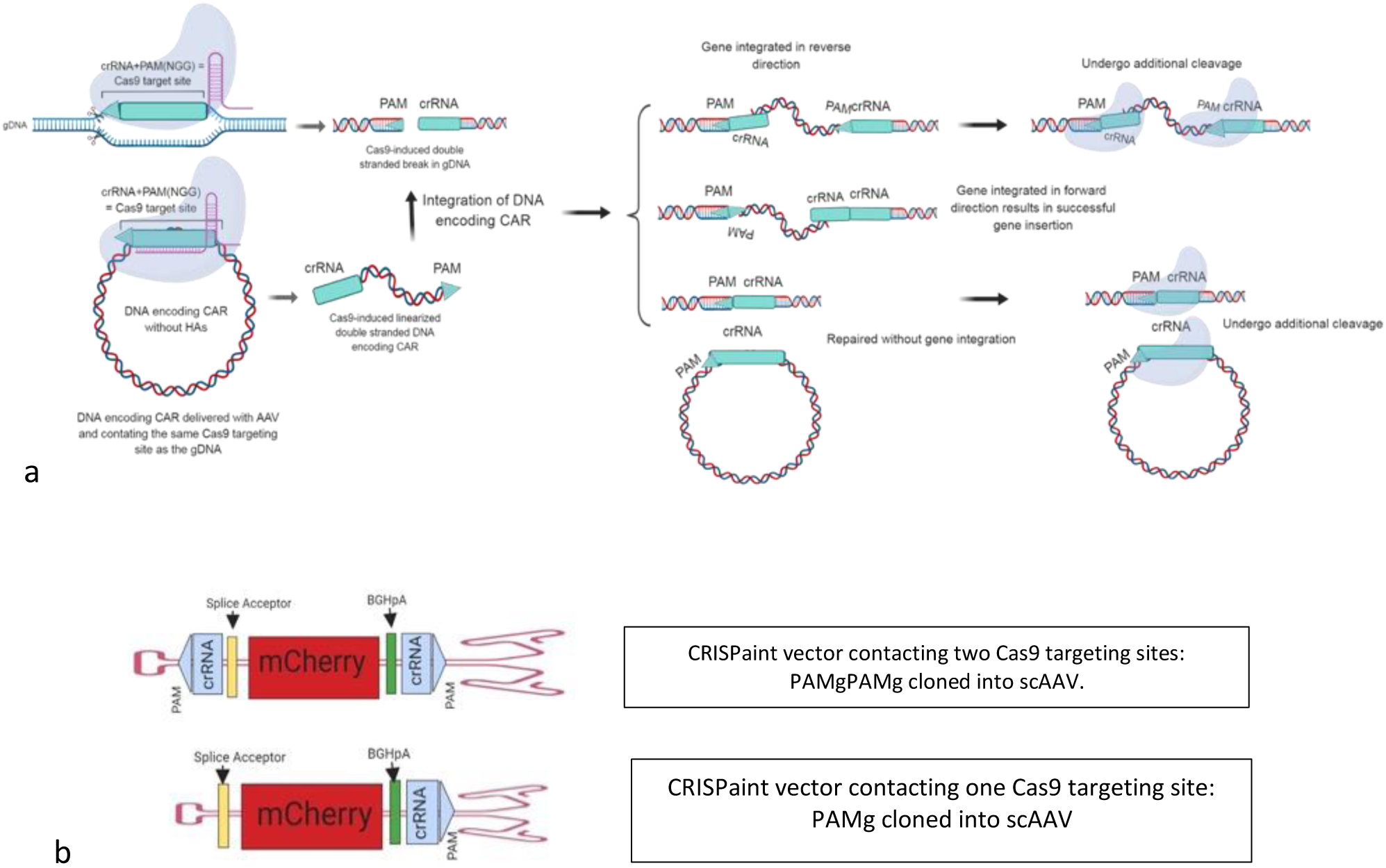
Gene integeration through CRISPaint approach does not require HA for flanking region of Cas9-targeting site. For non-homologues gene insertion through CRISPaint approach **(a)**, we included single (PAMg) or double (PAMgPAMg) AAVS1 targeting sites (crRNA+PAM sequences) within ITRs. The CRISPaint helps to overcome the the time-consuming process of designing HAs for HR-gene insertion **(b)**.

**Figure 10.**
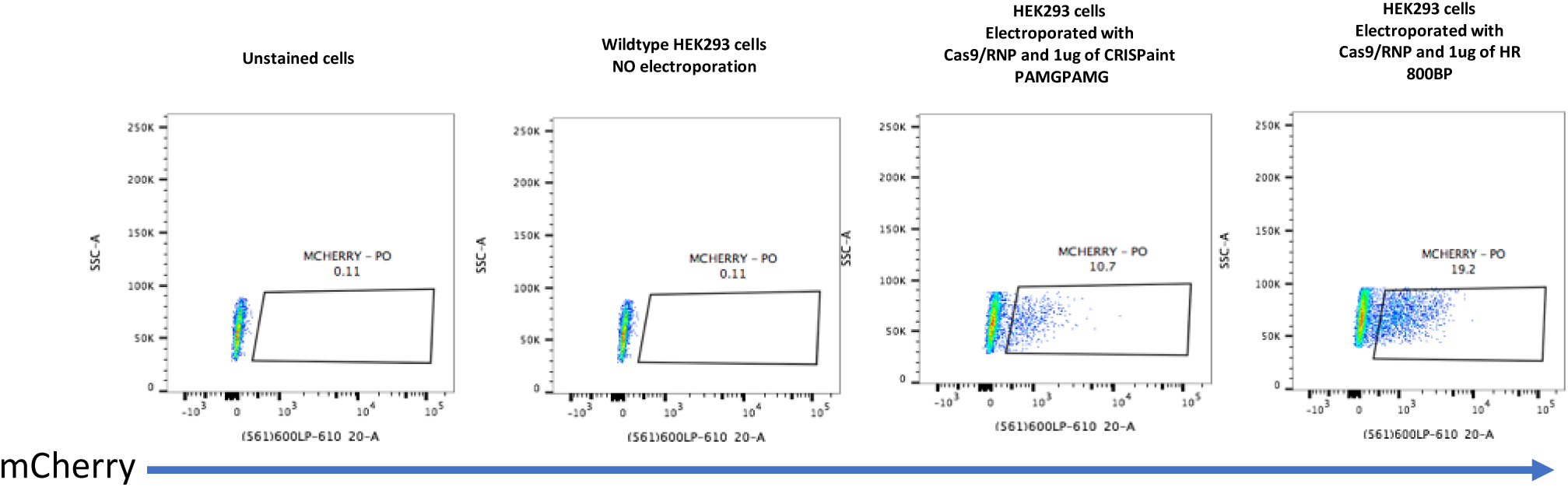
Flow cytometry results showed successful transgene expression in HEK293 cells. Electroporation of 1ug of DNA-encoding mCherry into HEK293 cells to assay the accuracy of the vectors showed successful gene expression. Flow cytometry showed the stable expression of mCherry.

### Studying the NHEJ and HR pathways in the primary NK cells to determine optimal pathway for genome editing

CRISPaint and HR are regulated by enzymatic reactions. CRISPaint is a LIG4-dependent process, while other proteins such as BRCA1 and BRCA2 regulate HR^11^. Therefore, we analyzed expression levels of these genes in NK cells to evaluate which repair pathway would likely be more efficient in this cell type. RNA-seq analysis showed that the expanded NK cells have higher expression of BRCA1 and BRCA2 in comparison to naïve NK cells and there is no decrease in LIG4 level in these cells (Figure 11), providing potential optimal conditions for either HR or NHEJ-directed gene insertion through CRISPaint.

**Figure 11.**
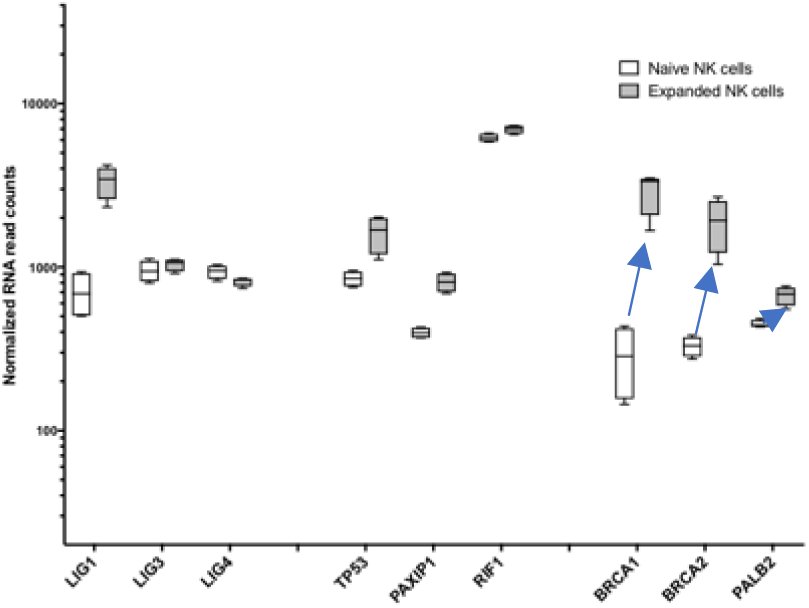
Studying the expression level of HR and CRISPaint regulating enzymes in naïve and expanded NK cells. HR related enzymes were upregulated in day-7 expanded NK cells in comparison to naïve NK cells. There is no difference of non-homologues related enzyme LIG4 between naïve and expanded NK cells. This data shows that day 7 expanded NK-cells have potential optimal condition for electroporation and transduction with AAV6 for directed gene insertion through both HR and CRISPaint.

### Combining Cas9/RNP and AAV6 to generate mCherry NK cells

A media change and resuspension at 5 × 10^5^ cells per ml was performed on day 6 of NK cell expansion one day prior to experimental manipulation. The NK cells were then electroporated with Cas9/RNP targeting AAVS1 on day 7 as described above. Thirty minutes after electroporation, live cells were collected and resuspended at 1 × 10^6^ cells per ml in media containing 100IU IL2 in a 24 well plate. For each transduction condition with ssAAV6 or scAAV6 to deliver HR or CRISPaint DNA encoding mCherry, we transduced 300K electroporated cells with 300K MOI or 150K MOI. Alternatively, to test the transduction efficiency of higher MOI of AAV6, we transduced 150,000 cells with 500K MOI of ssAAV6 delivering HR 800bp and scAAV6 delivering CRISPaint PAMgPAMg (Figure 12). Negative controls included NK cells that were not electroporated, were electroporated with Cas9/RNP but not AAV transduced, or were transduced with 300K MOI of AAV6 without electroporation of Cas9/RNP. The day after electroporation and transduction, we added 150ul of fresh media containing 100IU of IL2 to each well without changing the old media. The cells were kept in culture for 48 hours after electroporation and were then restimulated with K562 feeder cells at a ratio of 2:1 and kept in total volume of 1ml in 24 well plate.

**Figure 12.**
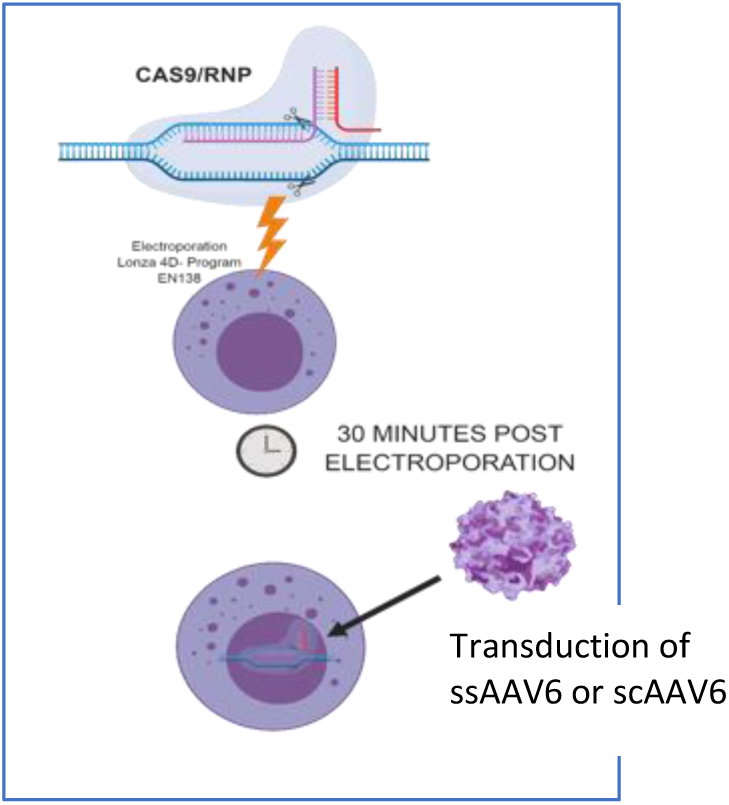
Combination of the electroporation of Cas9/RNP and transduction of AAV6 for gene insertion. 500K, 300K or 150K MOI of ssAAV6 or scAAV6 of HR and CRISPaint viruses delivering DNA-encoding mCherry were used to transduce the day 7 expanded NK cells which were electroporated with Cas9/RNP.

### Flow cytometry of CRISPR modified human NK cells shows successful integration of the mCherry gene

Two days after electroporation and transduction and before expansion, we performed flow cytometry to assess mCherry expression and viability (GhostRed 780 dye). As an overall, NK cells transduced with AAV6 delivering HR vectors had higher knock-in efficiency than CRISPaint. Furthermore, scAAV6 showed significantly better gene insertion. We did not observe mCherry expression in control NK cells. Almost 20% of NK cells in the experimental conditions in which HR vectors were used to transduce cells with the ssAAV6 delivering DNA-encoding mCherry with 800bp of HA, were mCherry positive. For NK cells which were transduced with 300K MOI of scAAV6 delivering CRISPaint PAMg or 300K MOI and 500K MOI of PAMgPAMg we found up to 8% of mCherry positive cells. Importantly, the percentage of mCherry positive cells was significantly higher in the cells that were transduced with the scAAV6 HR vectors containing the shorter homology arms. These conditions included transduction at 300K MOI or 150K MOI of HR scAAV6 vectors with 30bp (19-20%), 300bp (80-85%), 500bp (75-85%), and 800bp (80-89%). For the cells with lower transduction efficiency (ssAAV6-HR-800bp, scAAV6-CRISPaint PAMgPAMg), we enriched the mCherry positive population to 85% by FACS sorting and continued expanding them for up to 20 days using irradiated mbIL21-expressing K562 feeder cells with no changes in the percentages of mCherry positive NK cells (figure 13-15). We repeated the flow analysis for the expanded mCherry positive NK cells up to 20 days post transduction and saw no significant changes in percentages of mCherry in the cells generated by a combination of Cas9/RNP and ssAAV6 or scAAV6. Although, we saw lower efficiency of gene integration using CRISPaint compared to HR-directed gene insertion, this method is still very attractive because it allows researchers to integrate genes of interest into a user-defined locus with no need for designing homology arms. Moreover, we saw better expansion in NK cells transduced with CRISPaint vectors (data not shown).

**Figure 13.**
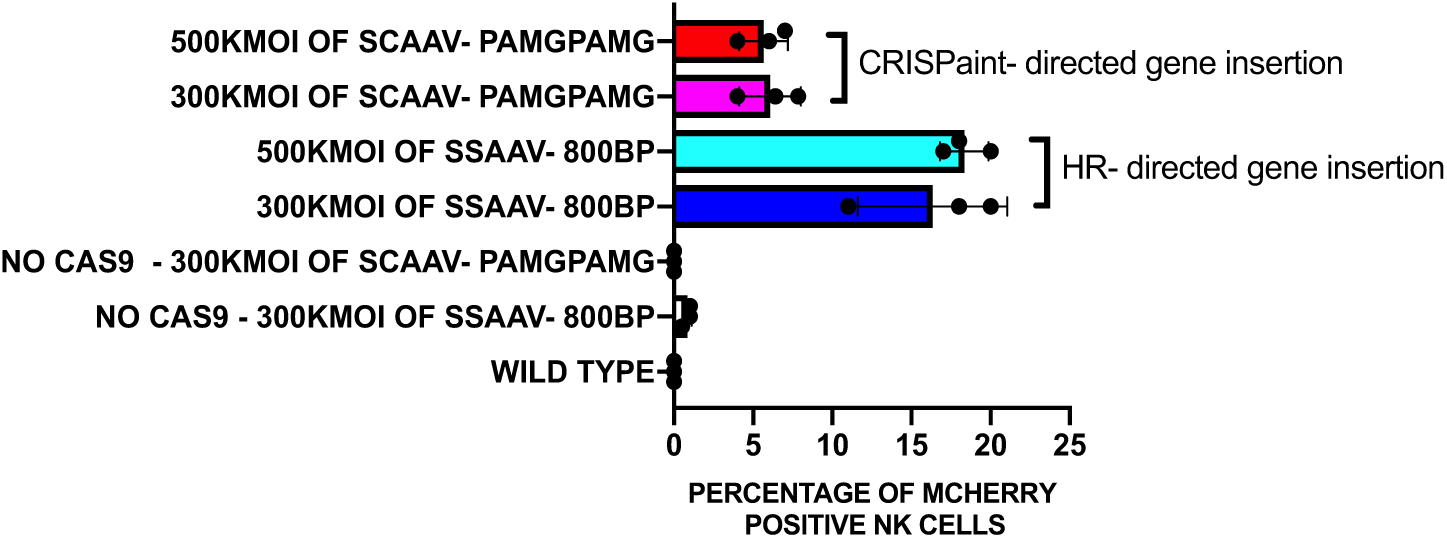
Flow cytometry results showed stable mCherry expression in CRISPR modified NK cells transuced with higher MOI of AAV6. We did not see any significant difference in the percentage of mCherry positive NK cells between the cells transduced with 500K MOI or 300K MOI of ssAAV delivering HR-800bp or scAAV delivering CRISPaint PAMgPAMg. Therefore, for the rest of the experiments we used 300K MOI and 150K MOI of AAV6.

**Figure 14.**
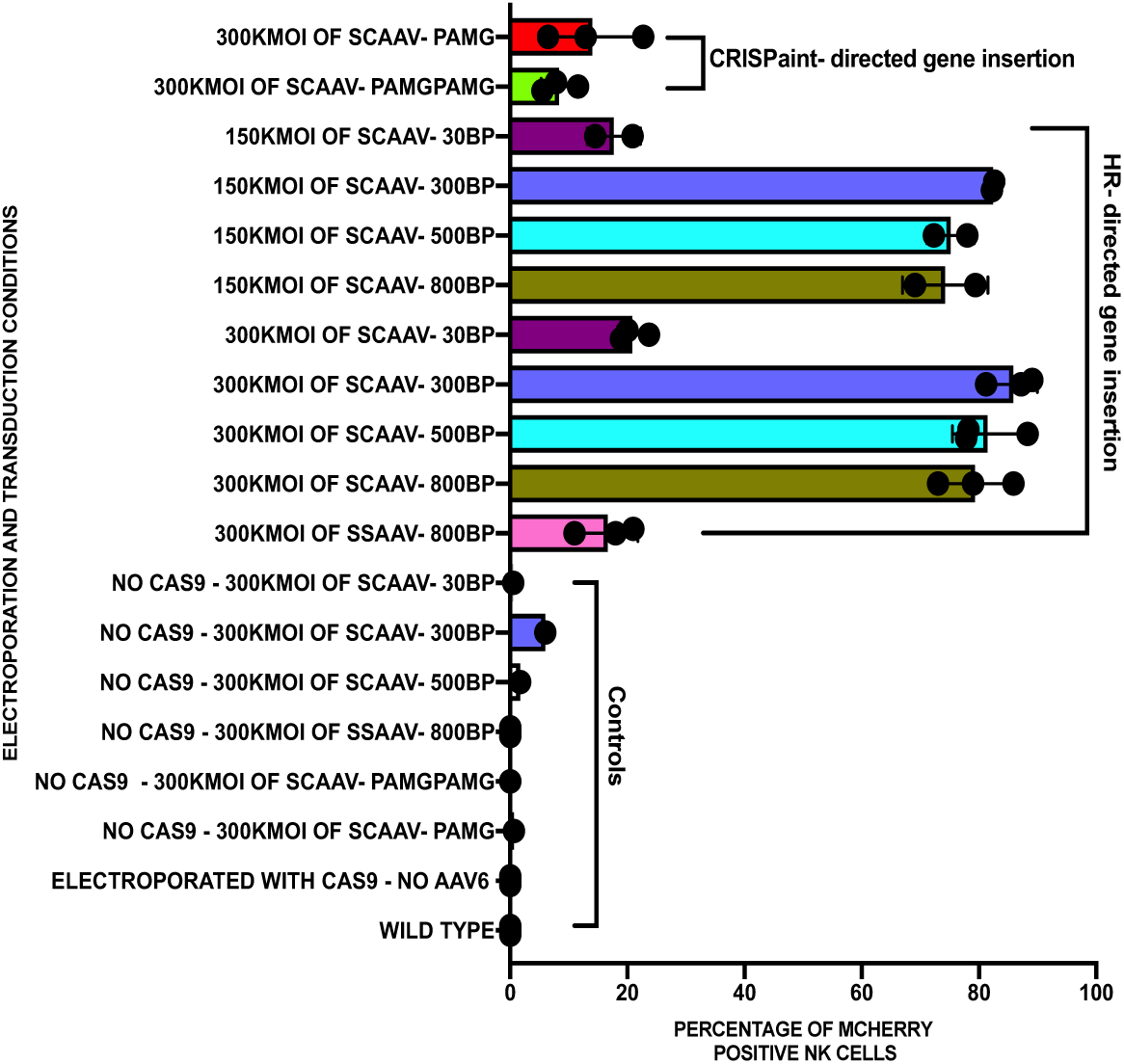
Flow cytometry results of CRIPSR modified primary NK cells demonstrated highly efficient transgene expression. We electroporated NK cells with Cas9/RNP and transduced with 300K MOI or 150K MOI of ssAAV or scAAV6 delivering HR or CRISPaint DNA template. Six days post CRISPR modification, flow cytometry results show that, delivering DNA encoding mCherry with minimum optimal homology arm lengths of 300bp for the flanking region of the Cas9-targeting site using scAAV6 results in highly efficient gene insertion in primary NK cells. larger HA including 500bp and 800-1000bp for smaller transgenes also can be utilized for gene insertion into NK cells. We also show that for the large transgenes 30bp of HA also can be used for gene insertion. Additionally, we demonstrate that CRISPaint is applicable approach for gene insertion into human primary NK cells.

**Figure 15.**
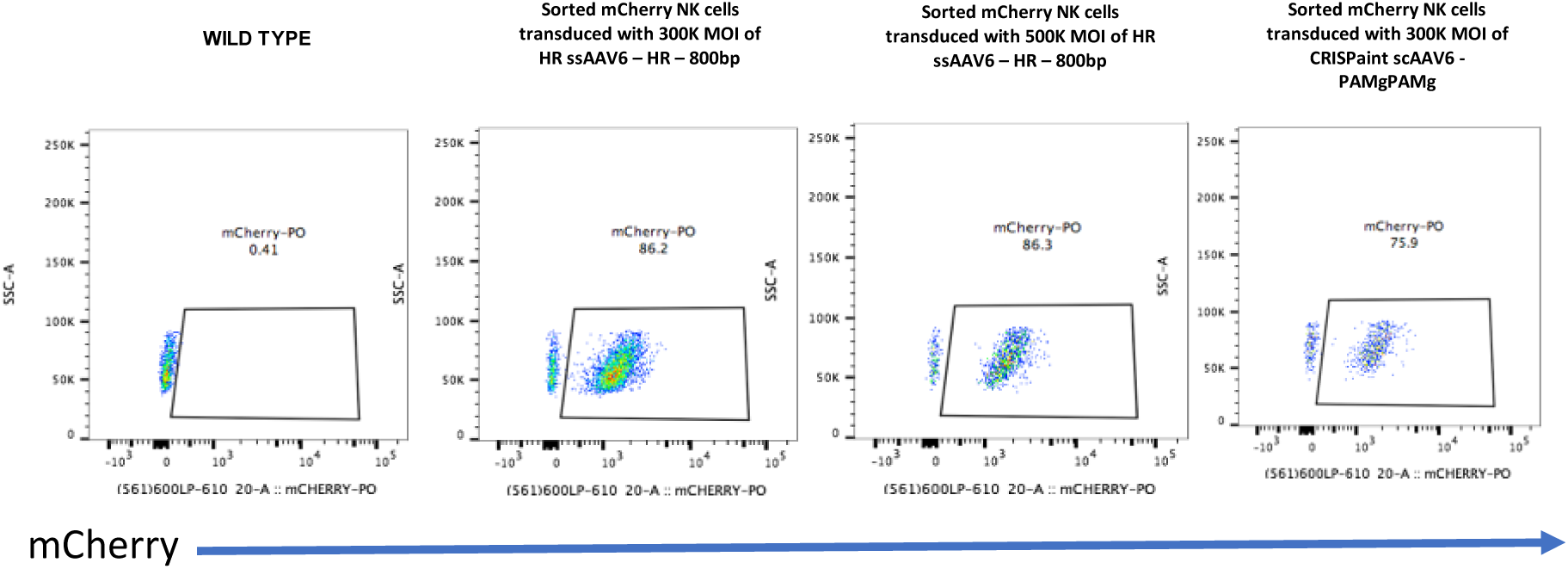
mCherry positive NK cells can be expanded on mb-IL21 expressing feeder cells. The mCherry NK cells which were generated with ssAAV6-HR-800bp and scAAV6-CRISPaint-PAMgPAMg and had lower efficiency were FACS sorted and grown for 20 days using feeder cells. The flow cytometry results 20 days post sorting showed the stable and enriched mCherry expression in CRISPR modified NK cells generated via both HR and CRISPaint.

### Combining Cas9/RNP with non-viral gene delivery causes cell death in primary human NK cells

In order to minimize the time and cost of virus production, non-viral gene integration has been used in T cells ^17^. We also tested this approach in NK cells by electroporating Cas9/RNP targeting AAVS1 with naked chemically-synthesized DNA encoding mCherry. The transgenes were generated in 2 forms, one as Megamer® Single-Stranded DNA Fragments from IDT with 80bp homology arms for the region flanking the Cas9-targeting site, and another with the same HR and CRISPaint DNAs that was cloned into the AAV backbone but were not packaged in any AAV capsids. Expanded NK cells were electroporated with Cas9/RNP targeting AAVS1 and 1 or 2ug of DNAs encoding mCherry in a total volume of 26ul. After 2 days, the cells with Megamer® were 100% dead, and only 10% of the cells which were electroporated with the HR and CRISPaint DNA had survived. We were able to further expand these cells, but less than 1% of the resulting cells were mCherry positive although the DNA was integrated at the site of DSB (data not shown). This indicates that AAV mediated NK cell modification is better tolerated and more efficient compared to naked DNA delivery.

## Discussion

Gene modification in primary human NK cells has always been challenging, here we report the first successful highly efficient site-directed gene integration into human primary NK cells using a combination of electroporation of Cas9/RNP and single stranded or self-complementary AAV6 gene delivery through HR and homology-independent gene insertion (CRISPaint). We show that a range of HAs from 30-1000bp can be used for gene insertion into the AAVS1 locus in NK cells, but that the optimal length is at 300bp. Additionally, we demonstrate that scAAV has significantly higher knock-in efficiency when combined with Cas9/RNP compared to ssAAV. We show that AAV6 is the best serotype for gene insertion into the genome of NK cells when it is combined with Cas9/RNP at 150K, 300K and 500K MOI, showing proof of concept for integrating genes into a defined locus in human primary NK cells. Our results also showed that the gene-modified NK cells could be subsequently expanded using irradiated feeders, enabling production of large numbers of gene-modified NK cells at clinical scale.

## Acknowledgments

We would like to thank Jonathan L. Schmid-Burgk from Feng Zhang’s lab at the Broad institute, Cambridge, MA for his help in reviewing the CRISPaint constructs..

